# Contribution of spatial heterogeneity in effective population sizes to the variance in pairwise measures of genetic differentiation

**DOI:** 10.1101/031633

**Authors:** Jérôme G. Prunier, Vincent Dubut, Lounès Chikhi, Simon Blanchet

**Affiliations:** Station d’Écologie Théorique et Expérimentale (UMR 5371), Centre National de la Recherche Scientifique (CNRS), Université Paul Sabatier (UPS), 2 route du CNRS, 09200 Moulis, France; Institut Méditerranéen de Biodiversité et d’Ecologie Marine et Continentale (UMR 7263), Aix-Marseille Université, CNRS, Institut de Recherche et du Développement, Avignon Université, Centre St-Charles, 3 place Victor Hugo, 13331 Marseille Cedex 3, France; Laboratoire Evolution & Diversité Biologique (UMR 5174), UPS, CNRS, Ecole Nationale de Formation Agronomique, 31062 Toulouse cedex 4, France; Instituto Gulbenkian de Ciência, Rua da Quinta Grande, n°6, 2780-156 Oeiras, Portugal.

**Keywords:** dispersal, evolutionary forces, genetic drift, landscape genetics, population genetics, variance partitioning

## Abstract

1. Pairwise measures of neutral genetic differentiation are supposed to contain information about past and on-going dispersal events and are thus often used as dependent variables in correlative analyses to elucidate how neutral genetic variation is affected by landscape connectivity. However, spatial heterogeneity in the intensity of genetic drift, stemming from variations in population sizes, may inflate variance in measures of genetic differentiation and lead to erroneous or incomplete interpretations in terms of connectivity. Here, we tested the efficiency of two distance-based metrics designed to capture the unique influence of spatial heterogeneity in local drift on genetic differentiation. These metrics are easily computed from estimates of effective population sizes or from environmental proxies for local carrying capacities, and allow us to introduce the hypothesis of Spatial-Heterogeneity-in-Effective-Population-Sizes (SHNe). SHNe can be tested in a way similar to isolation-by-distance or isolation-by-resistance within the classical landscape genetics hypothesis-testing framework.
2. We used simulations under various models of population structure to investigate the reliability of these metrics to quantify the unique contribution of SHNe in explaining patterns of genetic differentiation. We then applied these metrics to an empirical genetic dataset obtained for a freshwater fish (*Gobio occitaniae*).
3. Simulations showed that SHNe explained up to 60% of variance in genetic differentiation (measured as *Fst*) in the absence of gene flow, and up to 20% when migration rates were as high as 0.10. Furthermore, one of the two metrics was particularly robust to uncertainty in the estimation of effective population sizes (or proxies for carrying capacity). In the empirical dataset, the effect of SHNe on spatial patterns of *Fst* was five times higher than that of isolation-by-distance, uniquely contributing to 41% of variance in pairwise *Fst*. Taking the influence of SHNe into account also allowed decreasing the signal-to-noise ratio, and improving the upper estimate of effective dispersal distance.
4. We conclude that the use of SHNe metrics in landscape genetics will substantially improve the understanding of evolutionary drivers of genetic variation, providing substantial information as to the actual drivers of patterns of genetic differentiation in addition to traditional measures of Euclidean distance or landscape resistance.

## Introduction

The maintenance of effective dispersal capacities among demes is of tremendous importance for the viability of spatially structured populations (Wiens 1997). Given the technical challenges of directly monitoring individual movements, landscape genetics has emerged as an efficient way of assessing functional landscape connectivity (Tischendorf & Fahrig 2000), i.e. the influence of landscape configuration on effective dispersal between populations (i.e., gene flow; Barton & Bengtsson 1986), by combining methods from population genetics, landscape ecology and spatial statistics (Manel *et al*. 2003).

One of the main assets of landscape genetics is that it allows assessing functional landscape connectivity without the need for the quantitative inference of dispersal parameters (dispersal rate, dispersal distance or effective numbers of migrants; see Broquet & Petit 2009 for a review). Pairwise measures of neutral genetic differentiation (or “genetic distances”) such as F-statistics (e.g. *Fst*; Wright 1943) are supposed to contain information about past effective dispersal events (Slatkin 1987; Jaquiéry *et al*. 2011) and are thus considered a proxy for gene flow. The direct use of F-statistics as a proxy for gene flow ensues from the seminal work by Wright (1943) who showed that, under the specific assumptions of the island model, gene flow (the product of effective deme size *N_e_* and immigration rate *m*) between two populations could be derived from a measure of neutral genetic variance, following *Fst* = *1/(4Nm+1)*. Genetic distances are thus often used as dependent variables in correlative analyses to elucidate how neutral genetic variation is affected by landscape configuration (Storfer *et al*. 2007; Holderegger & Wagner 2008; Guillot *et al*. 2009). When carried out within a hypothesis-testing framework depicting the expected statistical relationships between landscape and neutral genetic data (Richardson *et al*. 2016), correlative analyses allow identifying the possible determinants of spatial genetic structures, thus providing a valuable way of assisting both landscape management and wildlife conservation (Segelbacher *et al*. 2010).

Depending on the complexity of the landscape, several competing hypotheses can be combined within the same analysis, such as *isolation-by-distance* (IBD) or *isolation-by-resistance* (IBR; Zeller *et al*. 2012). These hypotheses are formulated on the basis of how specific landscape features (coded as pairwise Euclidean or cost distances) are assumed to impact genetic differentiation and, by extension, gene flow. Isolation-by-distance is notably a baseline hypothesis in landscape genetics (Jenkins *et al*. 2010). In organisms whose dispersal ability is spatially constrained, the IBD hypothesis depicts the expected increase in genetic differentiation (and its variance) between populations as geographical distance increases. At a given spatial scale (e.g., Bradbury & Bentzen 2007), a significant positive correlation between Euclidean distances and genetic distances would give support to the tested hypothesis, suggesting that dispersal movements decrease as geographic distance increases (Slatkin 1993; but see Edelaar & Bolnick 2012).

However, it has to be emphasized that F-statistics are not estimates of gene flow *per se* but are primarily measures of the balance between genetic drift on the one hand, and migration (and mutations) on the other hand: high values of pairwise *Fst* indicate that genetic variation is mostly driven by drift, whereas low values indicate that genetic variation is mostly determined by dispersal, counterbalancing the effects of drift. Genetic drift is the evolutionary process of random fluctuations in allelic frequencies naturally occurring in all populations, whatever their size, though compounded in small ones (Allendorf 1986). If not fully compensated by gene flow, these random fluctuations ultimately lead to genetic differentiation, all the more quickly that at least one of the two considered demes is small. In other words, genetic distances may increase because of reduced landscape connectivity (and thus dispersal) between populations, but also because of spatial variations in population sizes (Jaquiéry *et al*. 2011; Richardson *et al*. 2016; see Appendix S1a for an illustration). Spatial heterogeneity in the intensity of genetic drift alone may thus be responsible for spurious relationships between measures of genetic differentiation and landscape predictors, erroneously providing support for alternative hypotheses such as IBD or IBR and possibly leading to counterproductive management and conservation measures. This risk of spurious conclusions is all the more important as the heterogeneity in the intensity of drift is not random but follows spatial patterns (such as upstream-downstream gradients in rivers or altitudinal gradient in mountains) that may in some cases be falsely captured by alternative landscape hypotheses (Appendix S1b). In any case, quantifying the contribution of spatial heterogeneity in local drift to the variance in genetic differentiation may provide crucial insights into the actual drivers of genetic structures.

In the same way that IBD or IBR hypotheses depict the expected contribution of landscape characteristics to the variance in genetic distances, we here propose an additional hypothesis relating the influence of spatial heterogeneity in local effective population sizes (*Ne*) over patterns of genetic differentiation: the Spatial-Heterogeneity-in-*Ne* (SHNe) hypothesis. This SHNe hypothesis, based on the computation of distance-based metrics from estimates of *Ne*, naturally falls within the hypothesis-testing framework classically used in landscape genetics, and is aimed at quantifying the contribution of heterogeneity in population sizes in shaping spatial patterns of genetic differentiation, thus providing further insight into acting evolutionary forces. Note that we did not intend to compute a measure of genetic differentiation “corrected” for *Ne* (as in Relethford 1996 or Jost 2008), but rather to provide metrics allowing a direct quantification of the amount of variance explained by SHNe.

In this study, we first described how estimates of *Ne* can be used to compute two distinct SHNe metrics stemming from the theoretical model of pure random genetic drift. We then investigated the influence of SHNe on the variance in *F_st_* values and on the behaviour of each metric in a simple two-deme situation. We then tested the ability and efficiency of each metric to account for heterogeneity in population sizes when they are directly computed from *Ne*, using simulations in various genetic models of population structure. Given the inherent difficulty in estimating *Ne* (Wang 2005), we used similar simulations to assess whether SHNe metrics were still efficient when computed from environmental proxies for local carrying capacities *K*, assuming that *K* is an imperfect proxy of *Ne*. We further assessed the efficiency of each metric in an empirical case study involving a freshwater fish species (*Gobio occitaniae*) using environmental estimates of population sizes. We finally discussed how and why these metrics should be recurrently used in landscape genetics studies.

## Materials and Methods

### DISTANCE-BASED METRICS OF SHNe

The *di* metric (for *distance based on the inverse*) was first proposed by Relethford (1991) and directly ensues from the classical formula depicting the expected loss of heterozygosity in an ideal Wright-Fisher population of constant size *N* over time but experiencing genetic drift (e.g. Hartl & Clark 2007, p. 122):

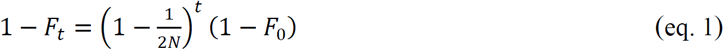

where *F_t_* is the fixation index at generation *t*. Extrapolated to a spatial context and with *F_0_* set to 0, the same equation can be used to depict the expected divergence of two subpopulations of size *Ni* relative to a founding population, in a situation where subpopulations are totally isolated, of constant size over time and with genetic drift being the only acting evolutionary force (Crow & Kimura 1970). After transformation (Relethford 1991), it can be shown that *F_st_* between populations 1 and 2 is proportional to *d_i_* (Appendix S2b), with:

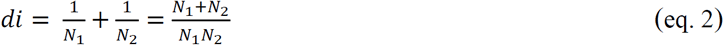

and *N_1_* and *N_2_* the (ideally effective) population sizes of populations 1 and 2, respectively.

The *dhm* metric (for *distance based on the harmonic mean*) was proposed by Serrouya *et al*. (2012) and also directly ensues from eq. 1. In the case of fluctuating population sizes over time, it can be shown that the effective size *Ne* of a population is the harmonic mean of census populations sizes *N* over time (Wright 1938; Hartl & Clark 2007). The harmonic mean weights smaller populations more heavily: in biological terms, it means that a single period of small population size (bottleneck) can result in a serious loss of heterozygosity. Extrapolated to a spatial context, the use of the harmonic mean entails that the smaller one of the two populations, the higher the pairwise genetic distance under a pure genetic drift model (Appendix S2a). We thus expect *Fst* between populations 1 and 2 to be proportional to *dhm*, with:

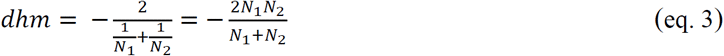

Note that we considered the opposite to the harmonic mean of *N* because the untransformed harmonic mean shows negative relationships with *F_st_* (Serrouya *et al*. 2012), and thus does not behave as a classical distance-based metric. Metrics *di* and *dhm* are inversely related and are thus expected to show different mathematical properties for the same combination of population sizes (Fig. 1b; Appendix S2c).

**Fig 1.**
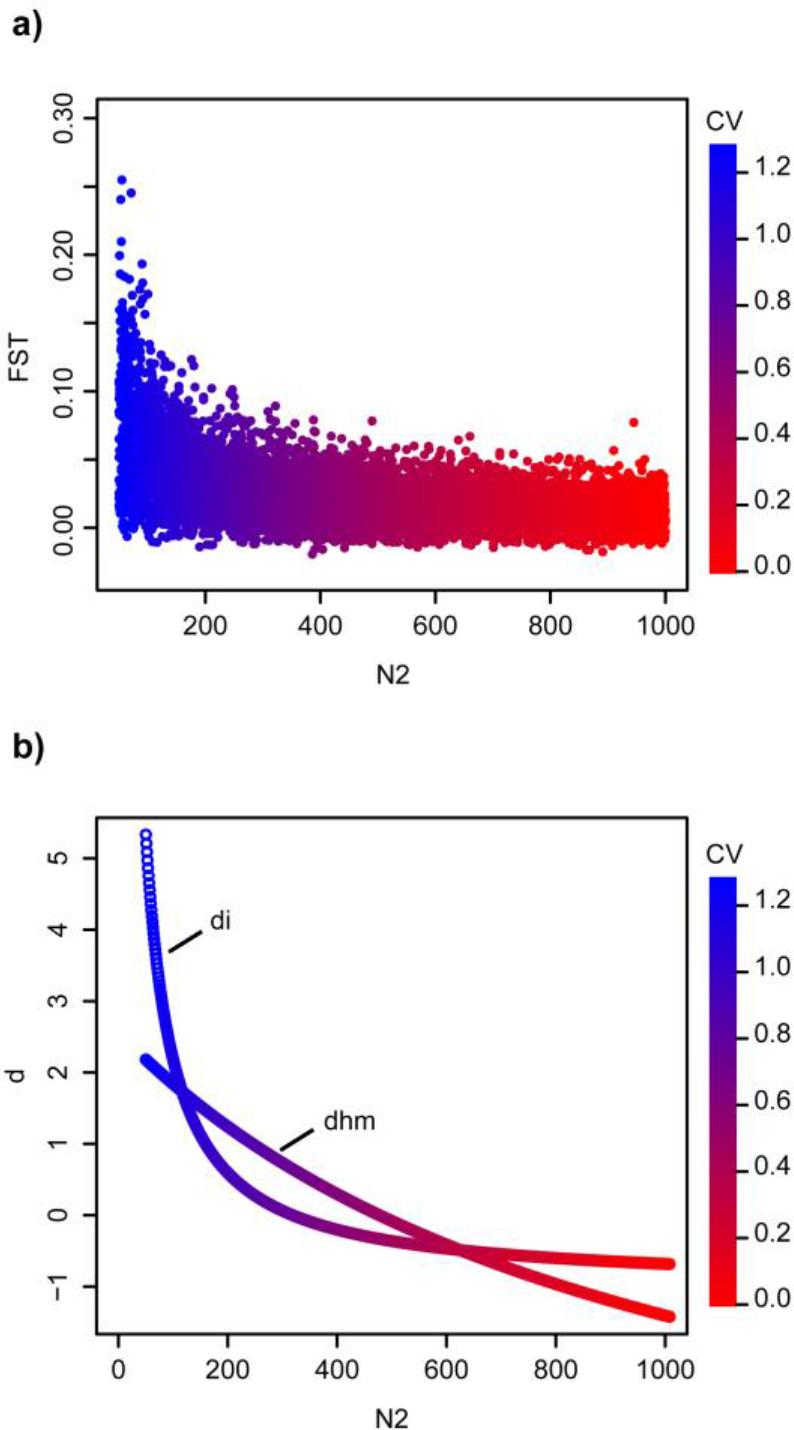
Behaviour of *Fst* (a) and each z-transformed SHNe metric (b) in a simple two-deme system as a function of effective population size *N_2_* (d: value of SHNe metrics; CV: Coefficient of variation).

Directly accounting for SHNe through the use of *Ne* is probably the most straightforward approach, but implies a major difficulty: estimating this demographic parameter, a task that may turn out to be tricky (Wang 2005). Alternatively, we thus propose to consider the use of environmental estimates of local carrying capacities (*K*) as a proxy for effective population sizes. Carrying capacity reflects the upper asymptote of the logistic growth curve of a population given the distribution and abundance of resources determined by local environmental conditions (Hanski 1994) and can be approximated using specific environmental variables such as habitat patch size or habitat quality (e.g. Raeymaekers *et al*. 2008).

### SIMULATED DATASETS

For all simulations, we used a computational pipeline including the programs ABCsampler (Wegmann *et al*. 2010), Simcoal2.1.2 (Laval & Excoffier 2004) and arlsumstat (Excoffier & Lischer 2010) to simulate and analyse microsatellite genetic datasets, with 15 independent loci following a stepwise mutation model and a unique mutation rate *µ* = 0.0005 (see Appendices S8 and S9 for results with *µ* = 0.01). Parameter values (symmetrical migration rates *m_i_* and local demes’ effective population sizes *Ne* (in number of haploid genotypes), hereafter only denoted as *N*) were picked from uniform distributions using ABCsampler and were then used as inputs in Simcoal2.1.2 to simulate genetic data based on a coalescent approach. In all simulations, a maximum of 30 haploid genotypes (that is, 15 diploid individuals) were sampled from each deme at the end of simulations and were used to compute pairwise *F_st_* among demes using arlsumstat. *di* and *dhm* metrics were computed on the basis of demes’ local carrying capacities *K*, with *K* = *N* + *α N*. The parameter represents the uncertainty in the estimates of effective population sizes through an environmental proxy such as habitat patch size. The estimates of *N* were considered as unbiased for α = 0 (since *K* = *N*) or uncertain for α ≠ 0. To control for heterogeneity in local genetic drift, effective population sizes *N_i_* were computed from a maximum population size *N_max_* following *N_i_* = *N_max_* - *γ N_max_*, with *N_max_* fixed at 1000 or randomly picked from a uniform distribution ranging from 100 to 1000 (see details below) and *γ* a correcting parameter randomly picked from a uniform distribution ranging from 0 to 0.95 so that *N_i_* ≤ *N_max_*. Levels of heterogeneity across effective population sizes were estimated using the coefficient of variation 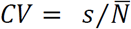, with *s* and 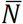 the standard deviation and the mean of effective population sizes, respectively. All statistical analyses were performed in R 3.1.2 (R Development Core Team 2014).

#### Influence of SHNe on the variance in *Fst*

We first investigated the influence of SHNe on the variance in *Fst* values and on the behaviour of each SHNe metric in a simple two-deme situation. We simulated 10^5^ genetic datasets with *m* fixed at 0.02 and set to 0. Effective population size for deme 1 (*N_1_*) was fixed at *N_max_* = 1000 (i.e. 500 diploid genotypes), while effective population size for deme 2 (*N_2_*) was computed from *N_1_* following *N*_2_ = *N*_1_ − *γ N*_1_. For each dataset, we computed pairwise *Fst, di* and *dhm* metrics, and *CV*. We then plotted *Fst* values as well as z-transformed *di* and *dhm* values against *N_2_*.

#### Contribution of SHNe metrics to the variance in *Fst*

We then assessed the contribution of each SHNe metric to the variance in *Fst* in four complex situations differing according to both the network structure and the migration model used for simulations (Appendix S3). We considered two different network structures: a one-dimensional 16-deme linear network and a two-dimensional 16-deme lattice network. We considered two distinct migration models: a spatially structured island model (Wright 1943) in which *m* decreases with Euclidean distance following an inverse-square function, and a spatially structured stepping-stone model (Kimura & Weiss 1964) where demes can only exchange migrants with adjacent demes. For each situation, 10,000 genetic datasets were simulated with set to 0, *m* randomly picked from a uniform distribution ranging from 0.0001 to 0.3 and *Nmax* randomly picked from a uniform distribution ranging from 100 to 1000. For each simulated dataset, we computed four pairwise matrices (*Fst, di, dhm* and Euclidean distances *mr*, the latter acting as a simple measure of inter-deme matrix resistance) and performed multiple regressions on distance matrices (MRDM; Smouse *et al*. 1986) between *Fst* and each SHNe metric, with *mr* as a unique covariate (*Fst* = *mr* + *di* or *Fst* = *mr* + *dhm*). All variables were z-transformed to standardize parameter estimates. Commonality analysis (a variance partitioning procedure assessing the reliability of model parameters in face of multicollinearity; Ray-Mukherjee *et al*. 2014; Prunier *et al*. 2015) was used to estimate the respective unique contribution (U) of each predictor to the variance in the dependent variable. The unique contribution is the part of the total variance in the dependent variable that is explained by the sole effect of the predictor being considered (*mr, di* or *dhm*). On the contrary, common contributions (shared variance among predictors) can help identify statistical suppression situations, responsible for artefactual distortions in parameter estimates (Paulhus *et al*. 2004; Prunier *et al*. 2017). Datasets were finally pooled according to their migration rate into 30 classes defined every 0.01 units (about 330 datasets per class). For each class, we computed the mean and the standard deviation of unique contributions of each predictor. A unique contribution was considered as negligible as soon as the dispersion around the mean included zero.

#### Minimum level of heterogeneity accounted for by SHNe metrics

To determine the minimum level of SHNe likely to affect pairwise *Fst*, we used the same approach as described above but with a constant migration rate (*m* = 0.02). For each simulated dataset, we additionally computed the coefficient of variation CV, and plotted the unique contribution of each predictor (*mr, di* or *dhm*) along with their respective standard deviation against CV.

#### Metrics measured from an environmental proxy

Finally, we investigated the influence of uncertainty in the estimation of *Ne* through an environmental proxy. We used the same approach as described above with *m* randomly picked from a uniform distribution ranging from 0.0001 to 0.3 and the parameter picked, independently for each population, from a uniform distribution ranging from -0.9 to 0.9 (Appendix S4).

### EMPIRICAL DATASET

As en empirical example, we considered neutral genetic data collected in the gudgeon (*Gobio occitaniae*), a small benthic freshwater fish. Fieldwork was conducted in accordance with French laws and with the approval of the Prefecture du Lot (the administrative region where the samples were collected). A total of 562 individuals were caught in 2011 by electro-fishing in 19 sampling sites scattered along the mainstream channel of the river Célé (Appendix S5). Pairwise *Fst* were computed between all pairs of sites using 11 microsatellite loci (Appendix S13). Details on laboratory procedures and acquisition of landscape data are provided in Appendix S6. The riparian distance among sites was used as a measure of matrix resistance *mr* among sites. We first used a simple Mantel test to assess significance of the relationship. We further investigated the observed pattern by using piecewise regression (Toms & Lesperance 2003) to identify the scale of migration-drift equilibrium, that is, the distance at which different linear relationships are observed. Estimations of breakpoints and slope parameters, as well as 95% confidence intervals, were performed using the R-package *segmented* (Muggeo 2008).

Given the difficulty in estimating effective population sizes in the wild (Wang 2005), we used a proxy for local carrying capacities to estimate *di* and *dhm*. The proxies we used were the width of the river at each sampling site and the estimated home-range size of each population (Raeymaekers *et al*. 2008). The home-range size of each deme was computed as the product of length and width of the river network (including tributaries) delimited by any downstream or upstream weir (see Blanchet *et al*. 2010 for a description of weirs in this river). This corresponded to the water area in which gudgeons were free to move. Matrices of pairwise *di* and *dhm* were then computed from these estimates and were independently confronted to the matrix of pairwise *Fst* using MRDM with 1000 permutations and with *mr* as a covariate. Commonality analyses were then used to disentangle the relative contribution of each predictor to the variance in pairwise *Fst*.

Finally, we plotted the residuals of the linear regression between *Fst* and the *di* metric (based on measures of river width) against *mr* and used a simple Mantel test to assess significance of the relationship. We further investigated the observed pattern by using piecewise regression to identify the distance threshold at which different linear relationships could be observed.

## Results

### SIMULATED DATASETS

#### Influence of SHNe on the variance in *Fst*

The aim of this first simulation study was to assess the behaviour of SHNe metrics in a very simple system in response to the variance in effective population sizes. Simulations were conducted so that any increase in heterogeneity (as measured by CV) was associated with a decrease in the size of one of the two populations (N_2_), the size of the other population (N_1_) being held constant. As expected, the increase in heterogeneity in *Ne* led to an increase in the variance in *Fst* (Fig.1a): *Fst* values ranged from 0 to 0.25 for a CV of 1.2 (N2 ≈ 5) while they did not exceed 0.05 for a CV of 0 (N2 ≈ 1000). The *dhm* metric showed a regular (though non-linear) increase with the increase in CV, while the *di* metric showed limited increase for CV ≤ 0.6, followed by a steep increase for CV > 0.6 (Fig.1b). When *Fst* and SHNe metrics were scaled to range from 0 to 1, SHNe metrics followed patterns similar to that observed for *Fst*, although *di* tends to better fit the general *Fst* pattern than *dhm* (Appendix S7).

#### Contribution of SHNe metrics to the variance in *Fst*

In the absence of uncertainty in the estimation of effective population sizes (α = 0), both *di* and *dhm* explained a non-negligible part of the total variance in pairwise *Fst* (i.e. from 5% to more than 60%; Fig. 2). Dispersion around the means further indicated that the unique contribution of SHNe metrics can be substantial, whatever the maximum population size in the system (*Nmax* ranging from 100 to 1000) and for migration rates as high as 0.135 (Fig. 2). For *m* < 0.135, the unique contributions of *di* and *dhm* were the lowest for a linear network with stepping-stone migration (Fig. 2a-b) and were the highest in a lattice network with spatially limited dispersal (Fig. 2g-h). In this latter case, *di* and *dhm* explained much more variance than the traditional covariate *mr* as soon as *m* < 0.05. For other genetic models, the unique contributions of *di* and *dhm* were roughly as high as (linear network with spatially limited migration; Fig. 2c-d) or slightly higher than (lattice network with stepping-stone dispersal; Fig. 2e-f) the unique contribution of *mr*. Overall, *di* and *dhm* behaved very similarly, although mean contribution as well as dispersion around the mean were slightly higher for *di* than for *dhm* in all situations.

**Fig 2.**
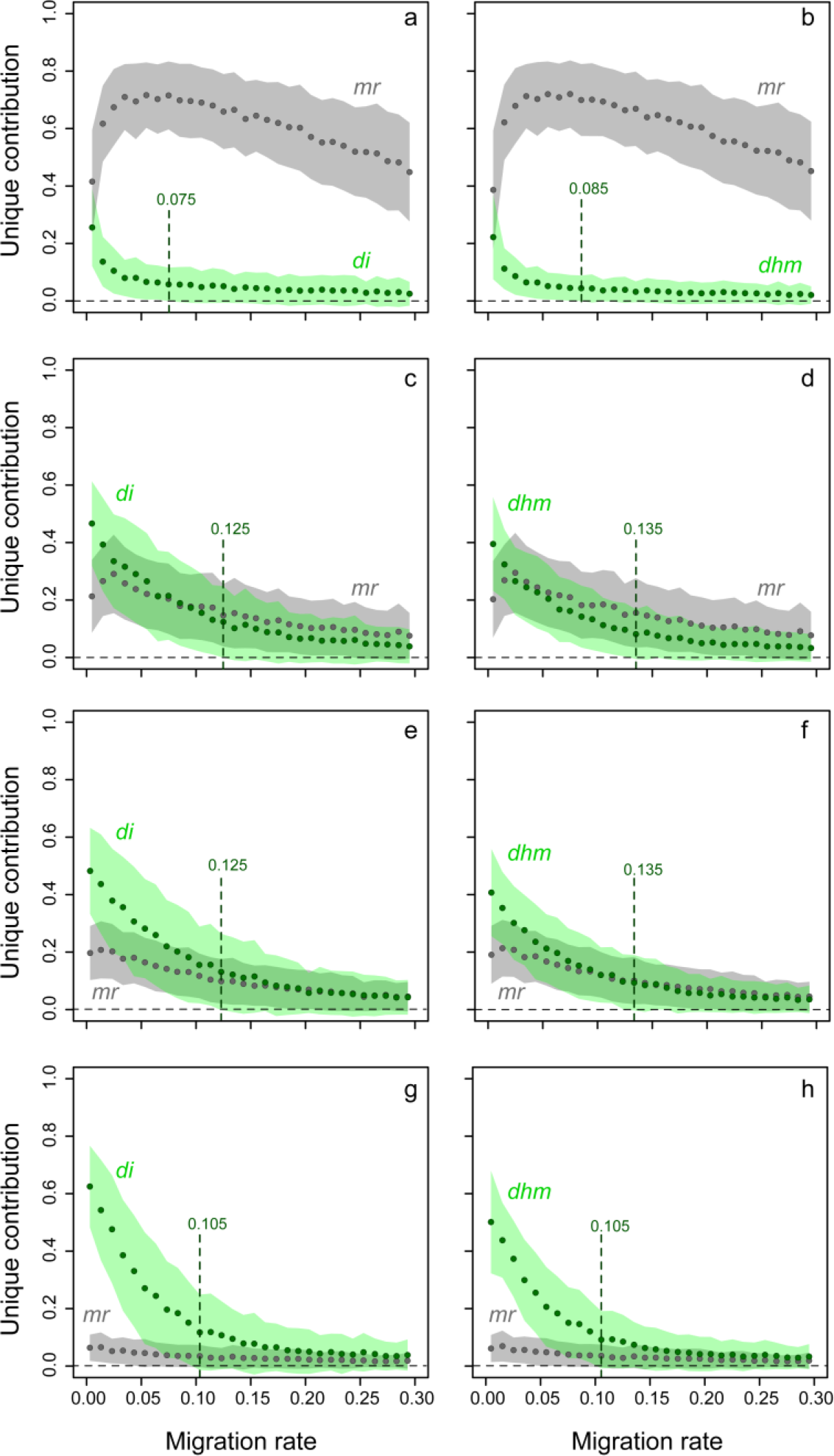
Unique contribution of metrics *mr* and *di* (left panels) or *mr* and *dhm* (right panels) as a function of the migration rate *m* for α = 0. Results are for a linear network with stepping-stone migration (a-b), a linear network with spatially limited dispersal (c-d), a lattice network with stepping-stone migration (e-f) and a lattice network with spatially limited dispersal (g-h). Circles represent the average unique contribution of each variable and coloured areas represent the dispersion of unique contributions around the mean, as defined by standard deviations. Vertical dashed lines indicate the migration rate *m* above which the unique contribution of SHNe metrics become negligible (lower bound for standard deviation < 0).

#### Minimum level of heterogeneity accounted for by SHNe metrics

When *m* was fixed at 0.02, the unique contribution of *dhm* and *di* were negligible when CV were below a value ranging from 0.23 (Fig. 3c-h) to 0.28 (linear network with stepping-stone migration; Fig. 3a-b). At maximum heterogeneity (CV ≈ 0.8), unique contributions ranged from 10 to 50% (Fig. 3). As previously, *di* and *dhm* behaved very similarly, although mean contribution as well as dispersion around the mean were slightly higher for *di* than for *dhm* in all situations. In most situations, the averaged model fit R² (the sum of unique and common contributions; Prunier *et al*. 2015) also increased with the increase of CV (e.g. from 10 to 60%; Fig. 3g-h). The observed decrease in the unique contribution of *mr* as the unique contribution of SHNe metrics increases (Fig. 3a-f) was not due to an increase in the common contribution of *mr* and SHNe metrics (data not shown), indicating that the relative support for alternative hypotheses such as IBD or IBR may be impaired in presence of SHNe.

**Fig 3.**
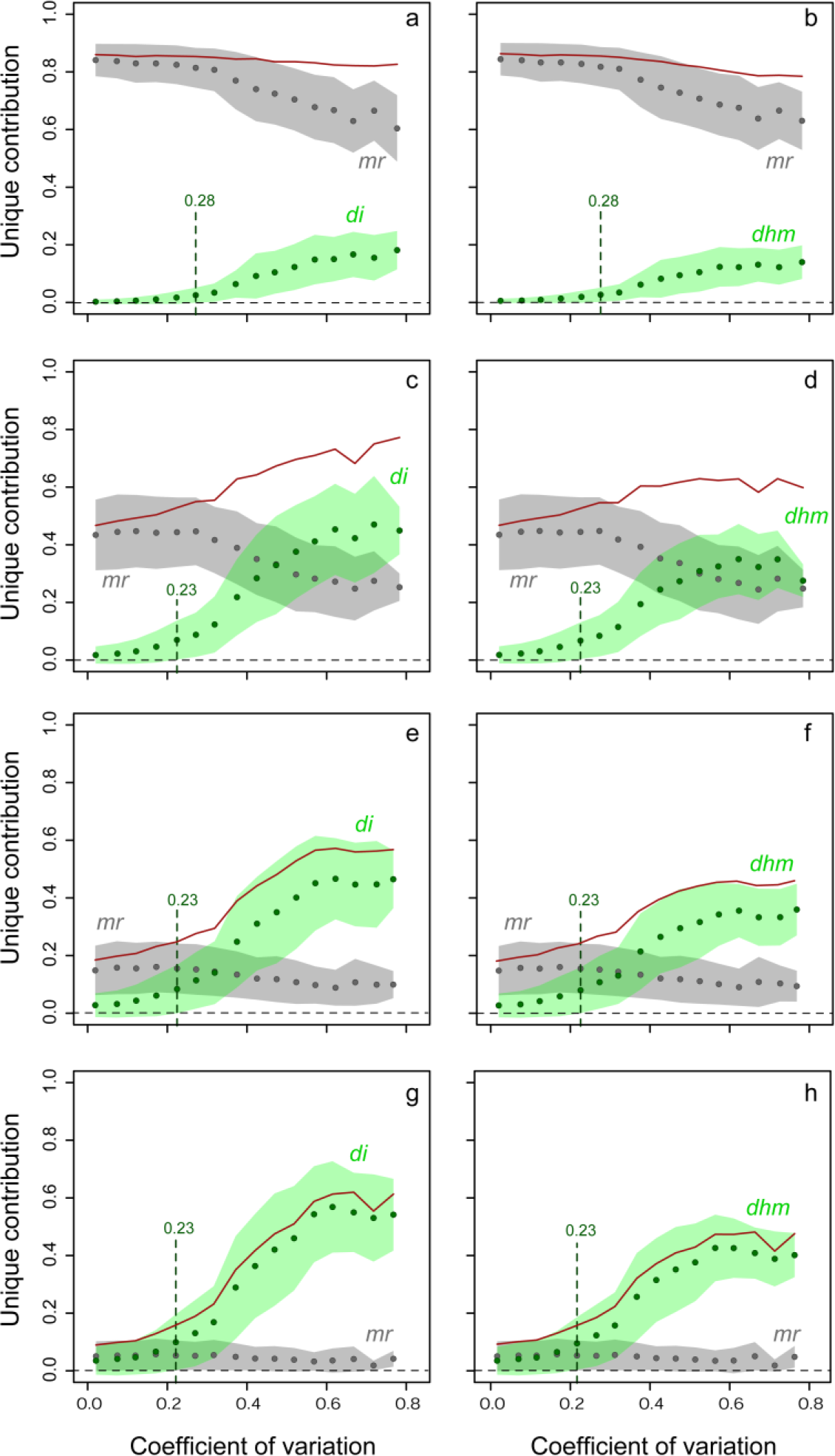
Averaged R^2^ (red) and unique contribution of metrics *mr* and *di* (left panels) or *mr* and *dhm* (right panels) as a function of heterogeneity in *Ne* (CV: coefficient of variation) for *m* = 0.02 and *α* = 0. See legend in Fig. 2 for other details.

#### Metrics measured from an environmental proxy

When uncertainty was included in the estimation of effective population sizes so as to mimic an environmental proxy for *K* (using α ϵ [-0.9, 0.9]), *mr* showed similar patterns to those in absence of uncertainty (Fig. 4). *SHNe* metrics behaved similarly to situations where true estimates of *N* were used, although unique contributions were systematically lower (but still up to 30% for *m* values lower than a threshold ranging from 0.025 to 0.105). Furthermore, and as previously, dispersion around the mean was generally noticeably larger for *dhm* than for *di*.

**Fig 4.**
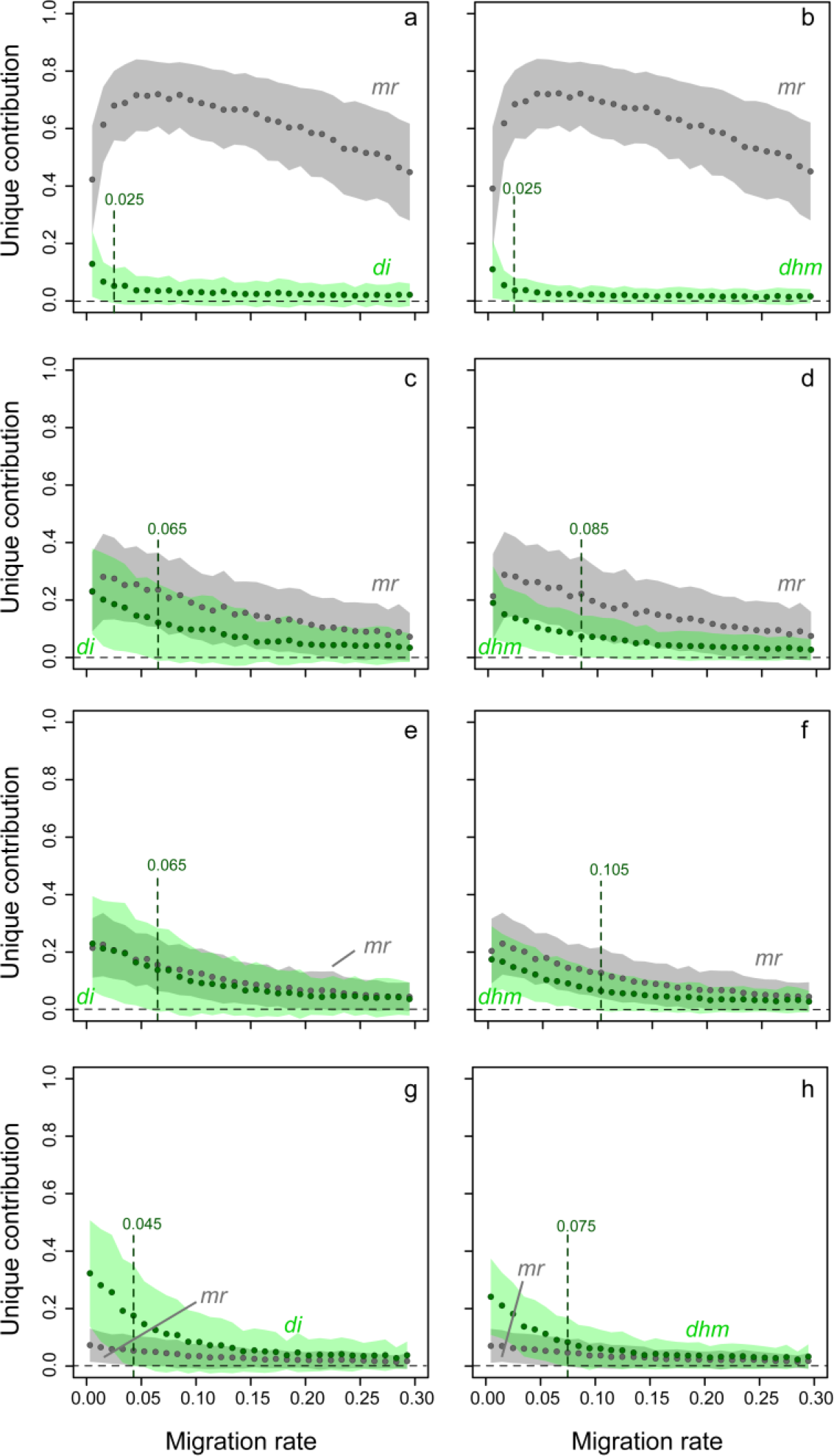
Unique contribution of metrics *mr* and *di* (left panels) or *mr* and *dhm* (right panels) as a function of the migration rate *m* for ranging from -0.9 to 0.9. See legend in Fig. 2 for other details.

### EMPIRICAL DATASET

The pattern of IBD in *G. occitaniae* was characterized by a slightly positive though insignificant relationship between *Fst* and *mr* (Table 1). Piecewise regression explained a slightly higher proportion of the variance in *Fst* (6.7%) than classical linear regression (2.9%). However, the upper bound of the confidence interval around the putative breakpoint (estimated at 74.8 ± 7.7 km) was located beyond spatial extent of the study.

**Table 1.**
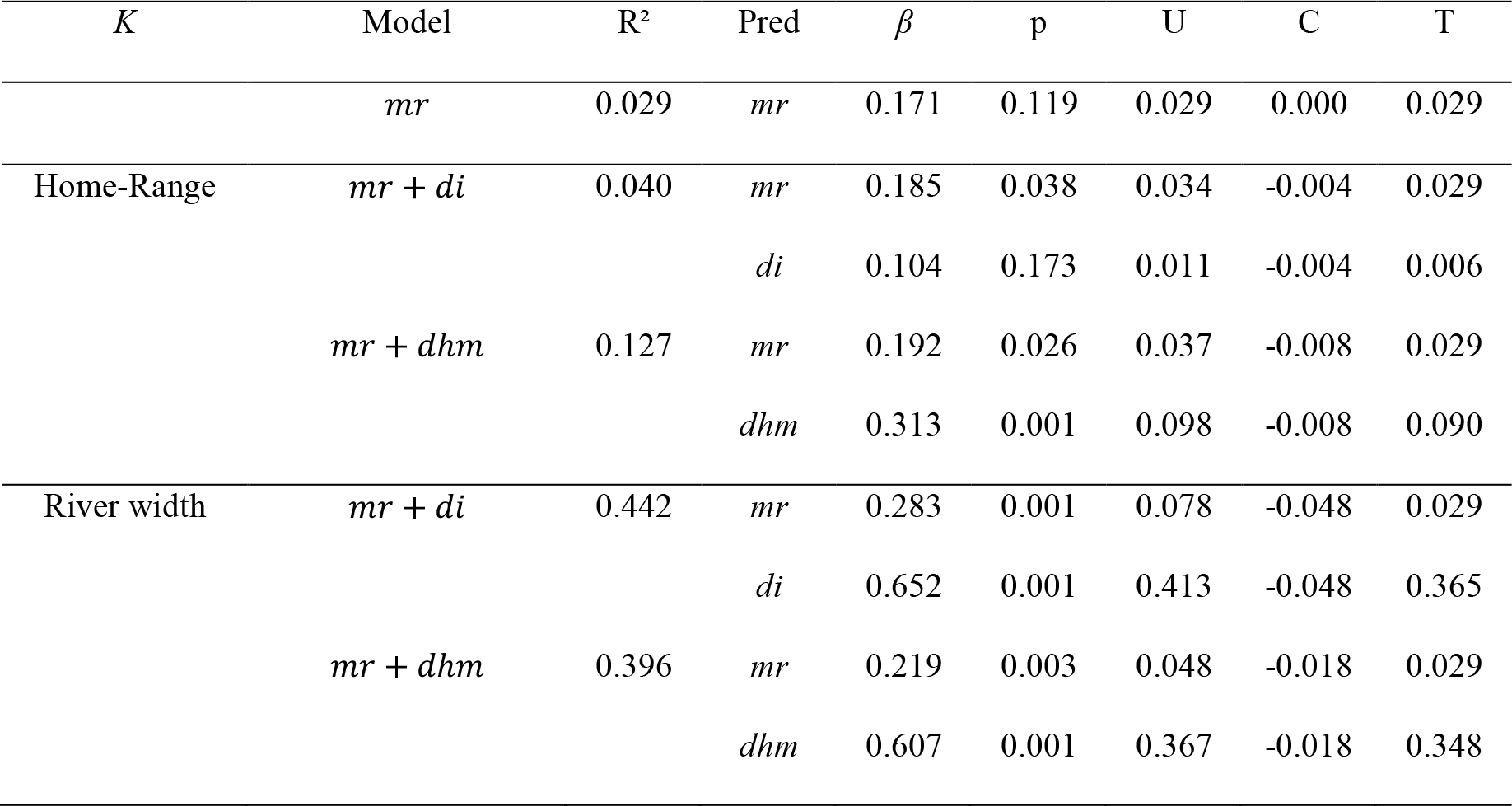
Results of Mantel test (univariate model), MRDM and commonality analyses (bivariate models) performed on empirical data. For each type of environmental proxy for carrying capacities (*K*) and each model (Model), the table provides the model fit index (R^2^) and, for each predictor (Pred), the standardized regression coefficients (beta weights *β*), the p-value (p) and finally unique (U), common (C) and total (T) contributions to the variance in the dependent variable.

When *K* values were estimated from river width or home-range sizes, coefficients of variation were respectively 0.432 and 0.927, suggesting high SHNe in this system. On the whole, model fit indices *R*^2^ were higher when *K* values were estimated from river width rather than from home-range sizes (Table 1), indicating that river width was a better proxy for carrying capacities than home-range size in this dataset. Whatever the proxy used for *K, mr* showed limited unique contribution to the variance in measures of genetic differentiation, with values ranging from 3.4 to 7.8% (Table 1). This variability in unique contributions of *mr* stemmed from collinearity with distance-based metrics of genetic drift, as revealed by common contributions C (Prunier *et al*. 2015): indeed, the highest unique contribution of *mr* (U = 7.8%) was also associated with the highest negative common contribution (C = - 4.8%), indicating statistical suppression, a situation responsible for an artificial boost in both the regression coefficient and its significance (Paulhus *et al*. 2004; Prunier *et al*. 2017). The observed variability in model fits (ranging from 4% to 44.2%) thus mostly ensued from the variability in SHNe metrics’ unique contributions to the variance in *Fst*. When *K* values were estimated from home-range sizes, the effect of *di* was not significant (unique contribution of 1.1%) whereas *dhm* uniquely accounted for 9.8% of variance in *Fst*. When *K* values were estimated from river width, the unique contribution of *di* and *dhm* strongly increased, reaching 41.3% and 36.7% respectively (Table 1).

When exploring the relationship between residuals of the linear regression between *Fst* and the *di* metric (based on measures of river width) and *mr*, piecewise regression explained a substantially higher proportion of the variance in *Fst* (23.8%) than linear regression (12.5%). The scatterplot showed an increase in residual values up to 8.9 ± 3.3 km and a clear-cut plateau beyond this threshold (Fig. 5b). This pattern suggests that, once the influence of SHNe is taken into account, the scale of IBD may be better inferred, with the divergence between populations located more than 8.9 km apart being mostly determined by drift.

**Fig 5.**
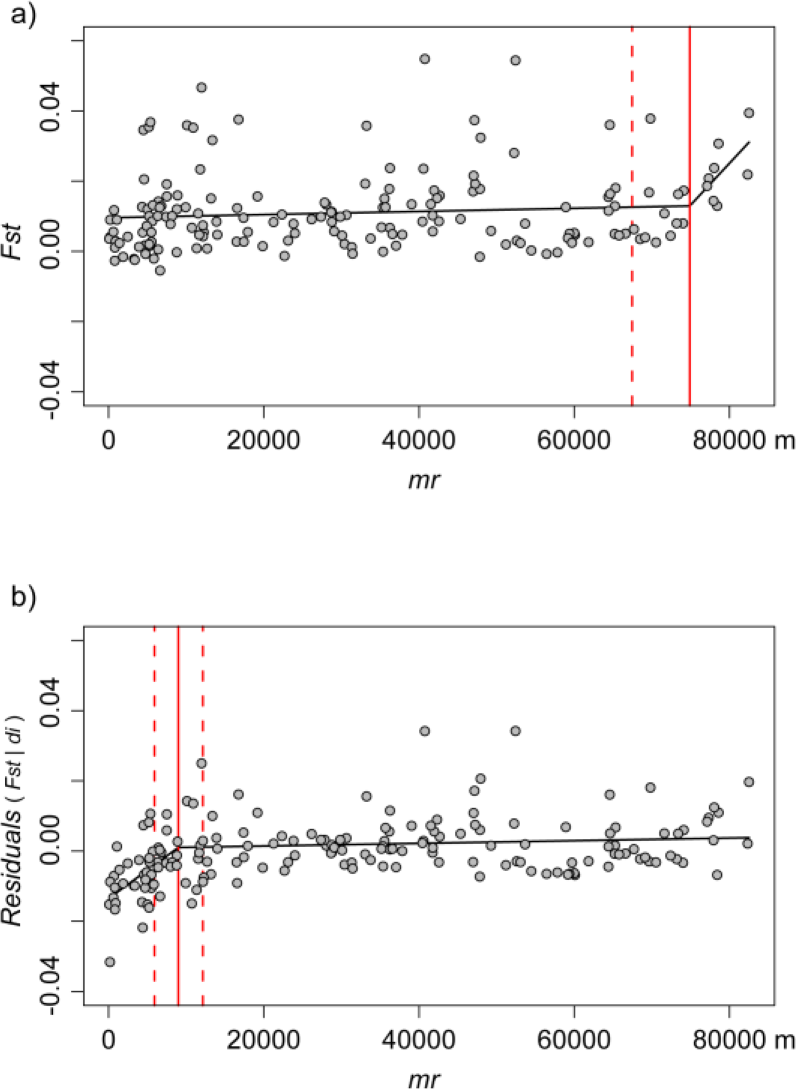
Scatterplot of pairwise *Fst* against (a) pairwise riparian distances (*mr*) and (b) residuals of the linear regression between *Fst* and the *di* metric (computed from river width) in the empirical dataset. Piecewise regression lines are in black. In red, solid vertical lines indicate the distance thresholds on either side of which different linear relationships can be observed, with 95% confidence intervals represented by dashed lines.

## Discussion

The possible influence of SHNe on the raw material of most landscape genetic studies, namely the variance in inter-deme measures of genetic differentiation, is rarely taken into consideration (but see Leblois *et al*. (2004) for an example), although it may lead to erroneous or incomplete interpretations of observed genetic patterns. Our study demonstrates that considering SHNe metrics (i.e., distance metrics measured from estimates of *Ne* or from environmental proxies for local carrying capacities) is a relevant approach to quantify the contribution of SHNe to the variance in pairwise measures of genetic differentiation, providing substantial information as to the actual drivers of observed patterns of genetic differentiation in addition to alternative hypotheses such as IBD. The proposed framework is based on a simple variance-partitioning procedure and does not require any complex parameterisation.

### COMPARISON OF SHNe METRICS

In a simple two-deme situation with constant migration rate, SHNe metrics exhibited patterns similar to *Fst*, thus properly rendering the influence of spatial heterogeneity in local drift on deme genetic differentiation. In more realistic scenarios, these metrics allowed quantifying the influence of SHNe on the variance in pairwise *Fst* from CV of about 25%. In other words, pairwise *Fst* in a set of three populations of mean size 100 and exchanging 2% of migrants at each generation (*m* = 0.02) may be affected by SHNe for differences in population sizes as low as 25 (e.g. *N* = 75, 100 and 125). The SHNe metrics explained up to 60% of variance in measures of genetic differentiation (and up to 80% for a higher mutation rate; see Appendix S8) at low migration rate, but still showed substantial contributions for migration rates as high as 0.135 (0.225 for a higher mutation rate). It is however noteworthy that the amounts of explained variance strikingly depended on the configuration of the network (1D *versus* 2D), the migration model (presence versus absence of long distance migration events) and the mutation rate (Appendices S8 and S9).

When applied to the *G. occitaniae* dataset, the use of *di* and *dhm* allowed explaining large amounts of variance in genetic differentiation when river width was used as a proxy for local carrying capacities. Only 2.9% of variance in measures of genetic differentiation was accounted for with *mr* as the only predictor. On the contrary, up to 44.2% of variance was explained when either *di* or *dhm* were used as additional explanatory variables. This suggests that the observed variance in measures of genetic differentiation was mostly driven by SHNe and much less by *IBD*. Interestingly, taking this effect into consideration through the use of residuals increased the scatterplot signal-to-noise ratio and revealed that genetic differentiation actually increased with riparian distance as long as populations were less than 8.9 km apart. This threshold corresponds to the scale of spatial autocorrelation in measures of genetic differentiation and may provide an upper estimate for effective dispersal distance (Anderson *et al*. 2010). This parameter is of particular interest as it may be used to inform conservation policies, for instance to support decisions in management of weirs or dams. Note however that taking the influence of SHNe into account in simulated datasets did not systematically lead to such an increase in signal-to-noise ratio (data not shown), probably because of variations in the spatial scale of simulated processes (Bradbury & Bentzen 2007). Further studies are hence needed to identify the specific conditions under which such improvement can be achieved: for instance, the upstream-downstream increase in carrying capacities encountered in river systems is responsible for a specific longitudinal pattern of SHNe (Appendix S1b) that may be more easily captured using this procedure than in the case of random SHNe (as simulated in this study).

When uncertainty was introduced in estimates of *Ne*, mimicking the use of an environmental proxy such as local carrying capacities, simulations showed that both *di* or *dhm* were still efficient at accounting for the influence of SHNe on the variance in *Fst*. However, *dhm* slightly outperformed *di* at intermediate migration rates (i.e., for *m* lower than 0.1) as *di*’s unique contribution to the variance in *Fst* showed higher dispersion around the mean than *dhm*. This trend was confirmed by the empirical dataset: when local carrying capacities were estimated from home-range sizes, *di* failed to detect any contribution of SHNe to the variance in genetic differentiation, whereas *dhm* -though less efficient than with river width as a proxy-still explained about 10% of variance. This difference stems from the inner characteristics of each metric. Given the use of a harmonic mean in its computation, *dhm* tends to increase as soon as one of the two demes shows a low to intermediate *Ne*. While *di* values show a rapid decrease as soon as one of the two demes shows an increase in *Ne*, the decrease in *dhm* values is smoother, thus still allowing the detection of the effect of SHNe despite higher uncertainty in the estimates of *Ne*. The *dhm* metric may actually be more robust when using environmental proxies for local carrying capacities and should therefore be preferred (or compared) to *di*. It is noteworthy that the two metrics can easily be combined in a single model and, provided collinearity patterns are inspected (Prunier *et al*. 2015), the best at fitting the dataset be selected according to its unique contribution (see Appendix S10 for an illustration).

### BIOLOGICALLY RELEVANT METRICS

Simulations indicated that the influence of SHNe was still perceptible for migration rates up to 0.15, irrespective of the model of population structure being considered (Figs 2 and 4). Interestingly, this range of values is higher than migration rates likely to be encountered in most natural systems. Indeed, summary statistics from 49 recent empirical studies that used BAYESASS (Wilson & Rannala 2003) to estimate interpatch migration rates (collected from a literature survey by Meirmans 2014; see Appendix S11 and Appendix S14 for details) indicated that the median value of average migration rates was 0.023, with more than 95% of studies showing average estimates lower than 0.1 (Appendix S12). For instance, the average estimate of migration rates in *G. occitaniae* in our empirical dataset was about 0.02 (unpublished data), as estimated using GENECLASS (Cornuet *et al*. 1999). These observations suggest that SHNe is likely to be an important driver of spatial genetic variation in many empirical datasets, considering that natural variability in deme sizes is most probably far from being an exception. Because of their ability to explain substantial additional amounts of variance in observed measures of genetic differentiation, we argue that considering the use of simple distance-based metrics such as *di* or *dhm* in future landscape genetic studies should thoroughly improve our understanding of observed spatial patterns of genetic variation.

### LIMITATIONS OF SHNe METRICS

Considering the difficulties in accurately estimating *Ne* from genetic data (Wang 2005), the use of alternative estimates of population size such as observed local densities (e.g. Joly *et al*. 2001; Blanchet *et al*. 2010) or habitat patch size (e.g. Verboom *et al*. 1991; Raeymaekers *et al*. 2008) to compute SHNe metrics is particularly appealing, but has yet to be considered with caution. The validity of such metrics indeed proceeds from the assumption that effective population sizes have remained constant over time (Appendix S2). This assumption theoretically limits the practical use of SHNe metrics to systems in which populations are not subject to abrupt changes in genetic drift. For populations having suffered from bottleneck events (Nei *et al*. 1975) or from founder effects (Ellstrand & Elam 1993), local environmental variables such as patch size may not properly mirror the actual effective population size, thus making SHNe metrics poor predictors of spatial patterns of genetic differentiation. In these situations, estimating effective population sizes from molecular data -although a delicate exercise-probably remains the best option (see Wang 2005 for a review synthesizing methods used to estimate *Ne*). More generally, integrating the demographic processes affecting *Ne* over time will be an important challenge to overcome so as to make landscape genetics an integrative discipline accounting for the complexity of spatially and temporally dynamic populations (Lowe & Allendorf 2010).

### CONCLUSION

Habitats modifications by humans have two components (Fischer & Lindenmayer 2007); one leading to a decrease in connectivity (fragmentation) and another leading to a decrease in habitat and resource availability (habitat loss and degradation). By reducing the size of available habitats and by decreasing connectivity among habitats, humans are rapidly making the ground more and more fertile for genetic drift, and therefore spatial heterogeneity in local drift, to become an increasingly influential evolutionary process. As the combined use of SHNe metrics and classical (IBD, IBR, etc.) landscape predictors in regression commonality analyses may substantially improve our understanding of how each process respectively contributes to observed spatial patterns of genetic variation, we believe that the time is ripe to systematically quantify the influence of SHNe on the spatial genetic structure of wild populations, in order to identify cases where genetic differentiation is actually increasing as a consequence of spatial heterogeneity in resource availability.

## Acknowledgements

We thank G. Loot, I. Paz-Vinas, O. Rey and C. Veyssière for their help on the field and in laboratory, as well as K. Saint-Pé for proofreading. We are grateful to Mark Beaumont for reading carefully a previous draft and providing comments that helped us clarify the manuscript significantly. We also thank the Office Nationale de l’Eau et des Milieux Aquatiques (ONEMA) for financial support, as well as two anonymous reviewers for their insightful comments and suggestions. Data used in this work were partly produced through the technical facilities of the Centre Méditerranéen Environnement Biodiversité. LC was partly funded by the LABEX (Laboratoire d’Excellence) entitled TULIP (vers une Théorie Unifiée des Interactions biotiques: rôle des Perturbations environnementales; ANR-10-LABX-41) and the LIA BEEG-B (Laboratoire International Associé - Bioinformatics, Ecology, Evolution, Genomics and Behaviour; CNRS). The authors declare no conflict of interest.

## Data accessibility

Simulated and empirical data: To be archived in Dryad

## Author Contributions

JGP and SB conceived and designed the study. SB and VD collected empirical data. VD generated molecular data. JGP performed the simulations and analysed simulated and empirical data. JGP, LC and SB interpreted the results and wrote the first draft of the paper and all authors contributed substantially to revisions.

### Supporting Information

**Appendix S1a.** An illustration of the influence of spatial heterogeneity in population sizes on the variance in *Fst*.

**Appendix S1b.** Neglecting SHNe may lead to spurious correlations.

**Appendix S2a.** The rationale behind the *dhm* metric

**Appendix S2b.** The rationale behind the *di* metric

**Appendix S2c.** Relationship between *dhm* and *di* metrics

**Appendix S3.** Network structures and migration models used in simulated datasets.

**Appendix S4.** Theoretical distribution of Pearson’s *r* correlation values between the effective population sizes *N* and local carrying capacities *K* for an uncertainty parameter ranging from -0.9 to 0.9.

**Appendix S5.** Main characteristics of the study area.

**Appendix S6.** Details on laboratory procedures and acquisition of landscape data in the empirical dataset.

**Appendix S7.** Behaviour of SHNe with respect to *Fst* values.

**Appendix S8.** Contribution of SHNe metrics to the variance in *Fst* for a mutation rate of 0.01.

**Appendix S9.** Contribution of SHNe metrics to the variance in *Fst* for a mutation rate of 0.01 and with uncertainty in the estimate of *Ne*.

**Appendix S10.** Empirical results from the full model, combining both *dhm* and *di* metrics.

**Appendix S11.** Selection criteria for empirical studies cited in the literature survey by P.G. Meirmans

**Appendix S12.** Distribution of average estimates of migration rates in 49 recent empirical studies.

**Appendix S13.** Summary statistics for the 11 microsatellite loci used in the empirical dataset.

**Appendix S14.** Summary statistics of the 49 retained empirical studies.

## References

Allendorf, F.W. (1986). Genetic drift and the loss of alleles versus heterozygosity. Zoo Biology, 5, 181–190.

Anderson, C.D., Epperson, B.K., Fortin, M.-J., Holderegger, R., James, P.M.A., Rosenberg, M.S., Scribner, K.T. & Spear, S. (2010). Considering spatial and temporal scale in landscape-genetic studies of gene flow. Molecular Ecology, 19, 3565–3575.

Barton, N. & Bengtsson, B.O. (1986). The barrier to genetic exchange between hybridising populations. Heredity, 57, 357–376.

Blanchet, S., Rey, O., Etienne, R., Lek, S. & Loot, G. (2010). Species-specific responses to landscape fragmentation: implications for management strategies. Evolutionary Applications, 3, 291–304.

Bradbury, I.R. & Bentzen, P. (2007). Non-linear genetic isolation by distance: Implications for dispersal estimation in anadromous and marine fish populations. ResearchGate, 340, 245–257.

Broquet, T. & Petit, E.J. (2009). Molecular Estimation of Dispersal for Ecology and Population Genetics. Annual Review of Ecology, Evolution, and Systematics, 40, 193–216.

Cornuet, J.-M., Piry, S., Luikart, G., Estoup, A. & Solignac, M. (1999). New methods employing multilocus genotypes to select or exclude populations as origins of individuals. Genetics, 153, 1989–2000.

Crow, J.F. & Kimura, M. (1970). An introduction to population genetics theory. xiv+591 pp.

Edelaar, P. & Bolnick, D.I. (2012). Non-random gene flow: an underappreciated force in evolution and ecology. Trends in Ecology & Evolution, 27, 659–665.

Ellstrand, N.C. & Elam, D.R. (1993). Population genetic consequences of small population-size - Implications for plant conservation. Annual Review of Ecology and Systematics, 24, 217–242.

Excoffier, L. & Lischer, H.E.L. (2010). Arlequin suite ver 3.5: a new series of programs to perform population genetics analyses under Linux and Windows. Molecular Ecology Resources, 10, 564–567.

Fischer, J. & Lindenmayer, D.B. (2007). Landscape modification and habitat fragmentation: a synthesis. Global Ecology and Biogeography, 16, 265–280.

Guillot, G., Leblois, R., Coulon, A. & Frantz, A.C. (2009). Statistical methods in spatial genetics. Molecular Ecology, 18, 4734–4756.

Hanski, I. (1994). A Practical Model of Metapopulation Dynamics. The Journal of Animal Ecology, 63, 151.

Hartl, D.L. & Clark, A.G. (2007). Principles of population genetics, 4th edn. Sinauer Associates, Sunderland, Mass.

Holderegger, R. & Wagner, H.H. (2008). Landscape genetics. Bioscience, 58, 199–207.

Jaquiéry, J., Broquet, T., Hirzel, A.H., Yearsley, J. & Perrin, N. (2011). Inferring landscape effects on dispersal from genetic distances: how far can we go? Molecular Ecology, 20, 692–705.

Jenkins, D.G., Carey, M., Czerniewska, J., Fletcher, J., Hether, T., Jones, A., Knight, S., Knox, J., Long, T., Mannino, M., McGuire, M., Riffle, A., Segelsky, S., Shappell, L., Sterner, A., Strickler, T. & Tursi, R. (2010). A meta-analysis of isolation by distance: relic or reference standard for landscape genetics? Ecography, 33, 315–320.

Joly, P., Miaud, C., Lehmann, A. & Grolet, O. (2001). Habitat matrix effects on pond occupancy in newts. Conservation Biology, 15, 239–248.

Jost, L. (2008). *G* _ST_ and its relatives do not measure differentiation. Molecular Ecology, 17, 4015–4026.

Kimura, M. & Weiss, G.H. (1964). The stepping stone model of population structure and the decrease of genetic correlation with distance. Genetics, 49, 561–576.

Laval, G. & Excoffier, L. (2004). SIMCOAL 2.0: a program to simulate genomic diversity over large recombining regions in a subdivided population with a complex history. Bioinformatics, 20, 2485–2487.

Leblois, R., Rousset, F. & Estoup, A. (2004). Influence of spatial and temporal heterogeneities on the estimation of demographic parameters in a continuous population using individual microsatellite data. Genetics, 166, 1081–1092.

Lowe, W.H. & Allendorf, F.W. (2010). What can genetics tell us about population connectivity? Molecular Ecology, 19, 3038–3051.

Manel, S., Schwartz, M.K., Luikart, G. & Taberlet, P. (2003). Landscape genetics: combining landscape ecology and population genetics. Trends in Ecology & Evolution, 18, 189–197.

Meirmans, P.G. (2014). Nonconvergence in Bayesian estimation of migration rates. Molecular Ecology Resources, 14, 726–733.

Muggeo, V.M.R. (2008). Segmented: an R package to fit regression models with broken-line relationships. R News, 1, 20–25.

Nei, M., Maruyama, T. & Chakraborty, R. (1975). The Bottleneck Effect and Genetic Variability in Populations. Evolution, 29, 1.

Paulhus, D.L., Robins, R.W., Trzesniewski, K.H. & Tracy, J.L. (2004). Two replicable suppressor situations in personality research. Multivariate Behavioral Research, 39, 303–328.

Pflüger, F.J. & Balkenhol, N. (2014). A plea for simultaneously considering matrix quality and local environmental conditions when analysing landscape impacts on effective dispersal. Molecular Ecology, 23, 2146–2156.

Prunier, J.G., Colyn, M., Legendre, X. & Flamand, M.-C. (2017). Regression commonality analyses on hierarchical genetic distances. Ecography.

Prunier, J.G., Colyn, M., Legendre, X., Nimon, K.F. & Flamand, M.C. (2015). Multicollinearity in spatial genetics: Separating the wheat from the chaff using commonality analyses. Molecular Ecology, 24, 263–283.

Raeymaekers, J.A.M., Maes, G.E., Geldof, S., Hontis, I., Nackaerts, K. & Volckaert, F.A.M. (2008). Modeling genetic connectivity in sticklebacks as a guideline for river restoration. Evolutionary Applications, 1, 475–488.

Ray-Mukherjee, J., Nimon, K., Mukherjee, S., Morris, D.W., Slotow, R. & Hamer, M. (2014). Using commonality analysis in multiple regressions: a tool to decompose regression effects in the face of multicollinearity. Methods in Ecology and Evolution, 5, 320–328.

Relethford, J.H. (1991). Genetic drift and anthropometric variation in ireland. Human Biology, 63, 155–165.

Relethford, J.H. (1996). Genetic drift can obscure population history: Problem and solution. Human Biology, 68, 29–44.

Richardson, J.L., Brady, S.P., Wang, I.J. & Spear, S.F. (2016). Navigating the pitfalls and promise of landscape genetics. Molecular Ecology, n/a-n/a.

Segelbacher, G., Cushman, S.A., Epperson, B.K., Fortin, M.-J., Francois, O., Hardy, O.J., Holderegger, R., Taberlet, P., Waits, L.P. & Manel, S. (2010). Applications of landscape genetics in conservation biology: concepts and challenges. Conservation Genetics, 11, 375–385.

Serrouya, R., Paetkau, D., McLellan, B.N., Boutin, S., Campbell, M. & Jenkins, D.A. (2012). Population size and major valleys explain microsatellite variation better than taxonomic units for caribou in western Canada. Molecular Ecology, 21, 2588–2601.

Slatkin, M. (1987). Gene flow and the geographic structure of natural populations. Science, 236, 787–792.

Slatkin, M. (1993). Isolation by distance in equilibrium and nonequilibrium populations. Evolution, 47, 264–279.

Smouse, P.E., Long, J.C. & Sokal, R.R. (1986). Multiple-regression and correlation extensions of the mantel test of matrix correspondence. Systematic Zoology, 35, 627–632.

Storfer, A., Murphy, M.A., Evans, J.S., Goldberg, C.S., Robinson, S., Spear, S.F., Dezzani, R., Delmelle, E., Vierling, L. & Waits, L.P. (2007). Putting the ‘landscape’in landscape genetics. Heredity, 98, 128–142.

Tischendorf, L. & Fahrig, L. (2000). On the usage and measurement of landscape connectivity. Oikos, 90, 7–19.

Toms, J.D. & Lesperance, M.L. (2003). Piecewise regression: a tool for identifying ecological thresholds. Ecology, 84, 2034–2041.

Verboom, J., Schotman, A., Opdam, P. & Metz, J.A.J. (1991). European nuthatch metapopulations in a fragmented agricultural landscape. Oikos, 61, 149–156.

Wang, J.L. (2005). Estimation of effective population sizes from data on genetic markers. Philosophical Transactions of the Royal Society B-Biological Sciences, 360, 1395–1409.

Wegmann, D., Leuenberger, C., Neuenschwander, S. & Excoffier, L. (2010). ABCtoolbox: a versatile toolkit for approximate Bayesian computations. BMC bioinformatics, 11, 116.

Wiens, J.A. (1997). 3 - Metapopulation Dynamics and Landscape Ecology. Metapopulation Biology (ed I.H.E. Gilpin), pp. 43–62. Academic Press, San Diego.

Wilson, G.A. & Rannala, B. (2003). Bayesian inference of recent migration rates using multilocus genotypes. Genetics, 163, 1177–1191.

Wright, S. (1943). Isolation by distance. Genetics, 28, 114.

Wright, S. (1938). Size of population and breeding structure in relation to evolution. Science, 87, 430–431.

Zeller, K.A., McGarigal, K. & Whiteley, A.R. (2012). Estimating landscape resistance to movement: a review. Landscape Ecology, 27, 777–797.

